# A Chinese hamster transcription start site atlas that enables targeted editing of CHO cells

**DOI:** 10.1101/2020.10.09.334045

**Authors:** Isaac Shamie, Sascha H. Duttke, Karen J. la Cour Karottki, Claudia Z. Han, Anders H. Hansen, Hooman Hefzi, Kai Xiong, Shangzhong Li, Sam Roth, Jenhan Tao, Gyun Min Lee, Christopher K. Glass, Helene Faustrup Kildegaard, Christopher Benner, Nathan E. Lewis

## Abstract

Chinese hamster ovary (CHO) cells, with their human-compatible glycosylation and high protein titers, are the most widely used cells for producing biopharmaceuticals. Engineering gene expression in CHO is key to improving drug quality and affordability. However, engineering gene expression or activating silent genes requires accurate annotation of the underlying regulatory elements and transcription start sites (TSSs). Unfortunately, most TSSs in the Chinese hamster genome were computationally predicted and are frequently inaccurate. Here, we revised TSS annotations for 15,308 Chinese hamster genes and 4,478 non-coding RNAs based on experimental data from CHO-K1 cells and 10 hamster tissues. The experimental realignment and discovery of TSSs now expose previously hidden motifs, such as the TATA box. We further demonstrate, by targeting the glycosyltransferase gene *Mgat3*, how accurate annotations readily facilitate activating silent genes by CRISPRa to obtain more human-like glycosylation. Together, we envision our annotation and data will provide a rich resource for the CHO community, improve genome engineering efforts and aid comparative and evolutionary studies.

## INTRODUCTION

Chinese hamster ovary (CHO) cells are the predominant mammalian system for large-scale production of clinical therapeutic proteins ^1^. They are valued for their high growth rate ^2^, ease of genetic manipulation and ability to properly fold, assemble, and produce complex post-translationally modified proteins that are not immunogenic in humans ^3^. As of 2018, 84% of FDA approved monoclonal antibodies were produced in CHO cells ^1^ and by sales, 10 of the top 15 drugs in the market are biopharmaceuticals and 7 are CHO-derived recombinant proteins ^1,4^. Optimizing CHO cells to increase production quantity and quality has been a priority for efforts to reduce the costs of biopharmaceuticals. Over the past few decades, these optimization efforts have progressed from engineering the media and bioreactors to transgene codon sequence and more recently, cell engineering and synthetic biology ^5,6^.

Genome sequencing efforts for the CHO and Chinese hamster ^7–9^ have been fundamental for studying and engineering CHO cells. In particular, they enabled systematic identification of genes associated with improved cell performance and product quality ^10–16^. Furthermore, the sequences enabled the implementation of CHO cell engineering using tools including transcription activator-like effector nucleases (TALENs, ^17^, RNA-directed DNA methylation (RdDM), ^18^, CRISPR-Cas9 ^19^) and others for genetic screens and the targeted inhibition or activation of genes ^6,20^. However, the Chinese hamster genome annotation remains far from complete, especially for the approximately 50% of genes that are silenced in CHO cells, including many needed for producing more human-like proteins ^21^. CRISPR activation (CRISPRa ^20^) and other genetic engineering methods could be instrumental to improve therapeutic protein production. However, to engineer gene expression, knowledge of the cis-regulatory elements and particularly a gene’s promoter and transcription start site (TSS) are critical. Recruitment of the RNA Polymerase II pre-initiation complex (RNAPII) by CRISPRa or blocking of the RNAPII by CRISPR inhibition (CRISPRi) or promoter editing ^22,23^, require knowledge of the polymerase’s native TSS. Unfortunately, the vast majority of cis-regulatory elements and transcript features in the Chinese hamster RefSeq annotation were predicted computationally. While previous work annotated 6,547 TSSs using steady-state 5’RNA ends by cap analysis gene expression (CAGE) in CHO cells ^24^, the data and annotation are not publicly available. Consequently, current inaccuracies in the annotation of the Chinese hamster genome and its TSSs present a major hurdle for targeted engineering of gene expression in CHO cells.

To remedy this issue, we experimentally re-annotated of the Chinese hamster transcriptome using multiple complementary experimental data types (GRO-seq ^25^, 5’GRO-seq ^26^, csRNA-seq ^27^, ribosomal RNA-depleted RNA-seq and ATAC-seq ^28^). To more comprehensively define regulatory elements in CHO cells, including for silenced genes, we captured nascent TSSs and open chromatin regions in CHO K1 cells, hamster bone marrow derived macrophages (BMDMs), and 10 hamster tissues from the original colony where CHO cells were generated ^29^. We observed TSSs in 15,308 protein-coding genes, updated TSS locations for 15,172 (70% of all genes) of them, and adjusted 8,921 (41% of observed genes) by more than 10 bps with the experimental TSSs (eTSSs). 30,760 promoter regions (TSSs within 150 bps) were observed across 15,308 Chinese hamster protein-coding genes and 4,478 promoter regions for non-coding RNAs. The accuracy of our new TSSs enable the analysis of regulatory DNA motifs, providing insights into the transcriptional regulatory pathways underlying tissue specificity. This resource sheds new light on gene regulation in the Chinese hamster, and will be valuable for optimizing the production of therapeutic recombinant proteins in CHO cells. To demonstrate the value of accurate TSSs, we identify a novel TSS 25 kb upstream of *Mgat3* (β-1,4-mannosyl-glycoprotein 4-β-N-acetylglucosaminyltransferase) that, when targeted with CRISPRa ^30^, increases the abundance of human-like bisecting N-acetylglucosamine on N-glycans ^5^ that are typically absent in CHO. Together, we envision our data and experimental TSS (eTSS) annotation for the Chinese hamster will provide a rich resource for the CHO community, improve engineering and manipulation of CHO cells, and enable comparative and evolutionary studies using the Chinese hamster.

## MATERIAL AND METHODS

### Sample preparation

Female Chinese hamsters (*Cricetulus griseus*) were generously provided by George Yerganian (Cytogen Research and Development, Inc) and housed at the University of California San Diego animal facility on a 12h/12h light/dark cycle with free access to normal chow food and water. All animal procedures were approved by the University of California San Diego Institutional Animal Care and Use Committee in accordance with University of California San Diego research guidelines for the care and use of laboratory animals. None of the used hamsters were subject to any previous procedures and all were used naively, without any previous exposure to drugs. Euthanized hamsters were quickly chilled in a wet ice/ethanol mixture (∼50/50), organs were isolated, placed into Trizol LS, flash frozen in liquid nitrogen and stored at −80C for later use. CHO-K1 cells were grown in F-K12 medium (GIBCO-Invitrogen, Carlsbad, CA, USA) at 37 °C with 5 % CO_2_.

### Bone marrow-derived macrophage (BMDM) culture

Hamster bone marrow-derived macrophages (BMDMs) were generated as detailed previously ^31^. Femur, tibia and iliac bones were flushed with DMEM high glucose (Corning), red blood cells were lysed, and cells cultured in DMEM high glucose (50%), 30% L929-cell conditioned laboratory-made media (as source of macrophage colony-stimulating factor (M-CSF)), 20% FBS (Omega Biosciences), 100 U/ml penicillin/streptomycin+L-glutamine (Gibco) and 2.5 μg/ml Amphotericin B (HyClone). After 4 days of differentiation, 16.7 ng/ml mouse M-CSF (Shenandoah Biotechnology) was added. After an additional 2 days of culture, non-adherent cells were washed off with room temperature DMEM to obtain a homogeneous population of adherent macrophages which were seeded for experimentation in culture-treated petri dishes overnight in DMEM containing 10% FBS, 100 U/ml penicillin/streptomycin+L-glutamine, 2.5 μg/ml Amphotericin B and 16.7 ng/ml M-CSF. For Kdo2-Lipid A (KLA), activation, macrophages were treated with 10 ng/mL KLA (Avanti Polar Lipids) for 1 hour.

### RNA-seq

RNA was extracted from organs that were homogenized in Trizol LS using an Omni Tissue homogenizer. After incubation at RT for 5 minutes, samples were spun at 21.000g for 3 minutes, supernatant transferred to a new tube and RNA extracted following manufacturer’s instructions. Strand-specific total RNA-seq libraries from ribosomal RNA-depleted RNA were prepared using the TruSeq Stranded Total RNA Library kit (Illumina) according to the manufacturer-supplied protocol. Libraries were sequenced 100 bp paired-end to a depth of 29.1-48.4 million reads on an Illumina HiSeq2500 instrument.

### csRNA-seq Protocol

Capped small RNA-sequencing was performed identically as described ^27^. Briefly, total RNA was size selected on 15% acrylamide, 7M UREA and 1x TBE gel (Invitrogen EC6885BOX), eluted and precipitated over night at −80°C. Given that the RIN of the tissue RNA was often as low as 2, essential input libraries were generated to facilitate accurate peak calling. csRNA libraries were twice cap selected prior to decapping, adapter ligation and sequencing. Input libraries were decapped prior to adapter ligation and sequencing to represent the whole repertoire of small RNAs with 3’-OH. Samples were quantified by Qbit (Invitrogen) and sequenced using the Illumina NextSeq 500 platform using 75 cycles single end.

### Global Run-On Nuclear Sequencing Protocol

Nuclei from hamster tissues were isolated as described ^32^. Hamster BMDM nuclei were isolated using hypotonic lysis [10 mM Tris-HCl pH 7.5, 2 mM MgCl_2_, 3 mM CaCl_2_; 0.1% IGEPAL] and flash frozen in GRO-freezing buffer [50 mM Tris-HCl pH 7.8, 5 mM MgCl_2_, 40% Glycerol]. 0.5-1 × 10^6^ BMDM nuclei were run-on with BrUTP-labelled NTPs as described ^33^ with 3x NRO buffer [15mM Tris-Cl pH 8.0, 7.5 mM MgCl_2_, 1.5 mM DTT, 450 mM KCl, 0.3 U/µl of SUPERase In, 1.5% Sarkosyl, 366 µM ATP, GTP (Roche) and Br-UTP (Sigma Aldrich) and 1.2 µM CTP (Roche, to limit run-on length to ∼40 nt)]. Reactions were stopped after five minutes by addition of 750 µl Trizol LS reagent (Invitrogen), vortexed for 5 minutes and RNA extracted and precipitated as described by the manufacturer.

### GRO-seq

RNA pellets were fully resuspended in 18 µl ddH_2_O + 0.05% Tween (dH2O+T). Subsequently, 2 µl fragmentation mix [100 mM ZnCl_2_, 10 mM Tris-HCl pH 7.5] was added and the samples were incubated at 70°C for 15 minutes. RNA fragmentation was stopped by placing samples on ice and addition of 2.5 µl 100 mM EDTA p.H. 8.0 and 500 μl cold GRO binding buffer [GRO-BB; 0.25×saline-sodium-phosphate-EDTA buffer (SSPE), 0.05% (vol/vol) Tween, 37.5 mM NaCl, 1 mM EDTA p.H. 8.0].

BrU enrichment was performed using a BrdU Antibody (Sigma B8434-200 µl Mouse monoclonal BU-33) coupled to Protein G (Dynal 1004D) beads. For each sample, 3x 20 µl of Protein G beads were washed twice in DPBS+0.05% Tween 20 (DPBS+T) and then the antibody coupled in a total volume of 1 ml DPBS+%T under gentle rotation. 1 µl of antibody was used per 8 µl of beads. After one hour at RT, the beads were collected using a magnet and the supernatant saved for future coupling reactions. Coupled beads were washed twice with GRO-BB prior to use. We pre-chill beads to 4 °C prior to the immunoprecipitation (IP) as traces of RNase may be observed. RNase activity was monitored using IDT’s RNaseAlert®. No crosslinking is required and worked best in our hands followed by DMP >> Bs3.

To each reaction of fragmented NRO-RNA with added GRO-BB, 20 μl equilibrated αBrdU beads were added and the samples slowly rotated at 4 °C for 80 minutes. Beads were subsequently collected on a magnet and the IP reaction washed twice with 500 μl GRO binding buffer before RNA was eluted by incubation of the beads in 500 μl Trizol LS (Thermo Fisher) under medium agitation for 5 minutes. 120 μl of TET [10 mM Tris-HCl pH 7.5, 1 mM EDTA p.H. 8.0, 0.05 % Tween20] was added to increase the supernatant and RNA extracted following manufacturer protocols.

For end-repair and decapping, RNA pellets were dissolved in 8 µl TET by vigorous vortexing, heated to 70°C for 2 minutes and placed on ice. After a quick spin, 22 µl Repair MM [3 µl 10x PNK buffer, 15.5 µl dH2O+T, 0.5 µl SUPERase-In RNase Inhibitor (10 U, Ambion), 2 µl PNK (20U, Enzymatics Y904L), 1 µl RppH (5U, NEB)] was added, mixed by flicking and incubated at 37°C for 1 hour. To phosphorylate the 5’end, 1 µl 10 mM ATP was subsequently added, and the reactions were incubated for another 45 minutes at 37°C (the high ATP concentration is advised to quench RppH activity). Following end repair, 2.5 µl 50 mM EDTA was added, reactions mixed and then heated to 70°C for 2 minutes before being placed on ice. A second BrdU enrichment was performed as detailed above.

For library preparation, RNA pellets were dissolved in 2.75 µl TET + 0.25 µl Illumina TruSeq 3’Adapter (10 mM), heated to 70°C for 2 minutes and placed on ice. 7 µl of 3’MM [4.75 µl 50% PEG8000, 1 µl 10x T4 RNA ligase buffer (NEB), 0.25 µl SUPERase-In, 1 µl T4 RNA Ligase 2 truncated (200U; NEB)] was added, mixed well by flicking and reactions incubated at 20°C for 1 hour. Reactions were diluted by addition of 10 µl TET + 2 µl 50 mM EDTA, heated to 70°C for 2 minutes, placed on ice and a third round of BdUTP enrichment was performed. RNA was transferred as a pellet to PCR strips during the 75% ethanol wash and dried. Samples were dissolved in 4 µl TET [10 mM Tris-HCl pH 7.5, 0.1 mM EDTA, 0.05% Tween 20] + 1 µl 10 mM reverse transcription (RT) primer. To anneal the RT-primer, the mixture was incubated at 75°C for 5 minutes, 37°C for 15 minutes and 25°C for ∼10 minutes. To ligate the 5’ Illumina TruSeq adapter, 10 µl 5’MM [1.5 µl ddH_2_O + 0.2% Tween20, 0.25 µl denatured 5’TruSeq adapter (10 mM), 1.5 µl 10x T4 RNA ligase buffer, 0.25 µl SUPERase-In, 0.2 µl 10 mM ATP, 5.8 µl 50% PEG8000, 0.5 µl T4 RNA ligase 1 (5U; NEB)] was added and reactions were incubated at 25°C for 1 hour. Reverse transcription was performed using Protoscript II (NEB) [4 µl 5x NEB FirstStrand buffer (NEB; E7421AA), 0.25 µl SUPERase-In, 0.75 µl Protoscript II (150U; NEB)] at 50°C for 1 hour. After adding 30 µl PCR MM [25 µl 2x LongAmp Taq 2X Master Mix (NEB), 0.2 µl 100 mM forward primer, 2.8 µl 5 M Betaine and 2 µl 10 mM individual barcoding primer], mixtures were amplified for 13 cycles. PCR reactions were cleaned up using 1.5 volumes of SpeedBeads™ (GE Healthcare) in 2.5 M NaCl/20%PEG8000 and libraries size selected on a PAGE/TBE gels to 150–225 bp. Gel slices were shredded by spinning through a 0.5 ml perforated PCR tube placed on top of a 1.5 ml tube. 150 µl Gel EB [0.1% LDS, 1M LiCl, 10 mM Tris-HCl pH 7.8] was added and the slurry incubated under agitation overnight. To purify the eluted DNA, 800 µl Zymogen ChIP DNA binding buffer (D5205) was added into the 1.5 ml tube containing the shredded gel slice and the Gel EB, mixed by pipetting and the slurry transferred to a ZymoMiniElute column. Samples were first spun at 1000g for 3 minutes, then 10k g for 30 seconds. Flow-through was removed, and samples washed with 200 µl Zymo WashBuffer (with EtOH). Gel remainders were removed by flicking and columns washed by addition on another 200 µl Zymo WashBuffer. Flow-through was removed, columns spun dry by centrifugation at 14k g for 1 minute and DNA eluted by addition of 20 µl pre-warmed Sequencing TET [10 mM Tris-HCl pH 8.0, 0.1 mM EDTA, 0.05% Tween 20]. Libraries were sequenced on an Illumina NextSeq 500 at using 75 cycles single end.

### 5’GRO-seq

RNA pellets were resuspended in 10 µl TET, heated to 70°C for 2 minutes and placed on ice. 10 µl of dephosporylation MM [2 µl 10x CutSmart, 6.75 µl dH2O+T, 1 µl Calf Intestinal alkaline Phosphatase (10 U; CIP, NEB) or quick CIP (10 U, NEB), 0.25 µl SUPERase-In (5U)] was added. Following incubation at 37°C for 45 minutes, 2 µl 50 mM EDTA was added, reactions mixed, heated to 70°C for 2 minutes and placed on ice. BrdU enrichment was performed as described for GRO-seq. RNA pellets were dissolved in 10 µl TET and a second round of dephosphorylation and BrdU enrichment was performed. Libraries were prepared as described in Hetzel et al. (2016). Briefly, libraries were done as described for GRO-seq (above) with exception of the 3’Adapter ligation step. Here, prior to 3’Adapter ligation, samples were dissolved in 3.75 µl TET heated to 70°C for 2 minutes and placed on ice. RNAs were decapped by addition of 6.25 µl RppH MM [1 µl 10x T4 RNA ligase buffer, 4 µl 50% PEG8000, 0.25 µl SUPERase-In, 1 µl RppH (5U)] and incubated at 37°C for 1 hour. Afterwards, to ligate the 3’ Illumina TruSeq adapter 10 µl of 3’MM was added [1 µl 10x T4 RNA ligase buffer, 6 µl 50% PEG8000, 1.5 µl ddH_2_O+T, 0.25 µl heat-denatured Illumina TruSeq 3’Adapter, 0.25 µl SUPERase-In, 1 µl T4 RNA Ligase 2 truncated K227Q (200U; NEB)] was added, mixed well by flicking and reactions incubated at 20°C for 1 hour. Reactions were diluted by addition of 10 µl TET + 2 µl 50 mM EDTA, heated to 70°C for 2 minutes, placed on ice and a third round of BrUTP enrichment was performed. 5’ adapter ligation, reverse transcription and library size selection were performed as described for GRO-seq. Samples were amplified for 14 cycles, size selected for 160–250 bp and sequenced on an Illumina NextSeq 500 at using 75 cycles single end.

### Assay for Transposase-Accessible Chromatin Sequencing (ATAC-Seq) Protocol

To approximately 150k nuclei in 22.5 µl GRO freezing buffer, isolated as described for GRO-seq above, 25 µl 2x DMF DNA Tagmentation buffer was added [66mM Tris-acetate(pH=7.8), 132 K-Acetate, 20mM Mg-Acetate, 32 % DMF], reaction mixed and 2.5 µl DNA Tagmentation Enzyme mix (Nextera DNA Library Preparation Kit, Illumina) added. Mixture was incubated at 37°C for 30 minutes and subsequently purified using the Zymogen ChIP DNA purification kit as described by the manufacturer. DNA was amplified using the Nextera Primer Ad1 and a unique Ad2.n barcoding primers using NEBNext High-Fidelity 2x PCR MM for 10 cycles. PCR reactions were purified using 1.5 volumes of SpeedBeads in 2.5M NaCl, 20% PEG8000, size selected for 140 – 240 bp, DNA eluted as described for GRO-seq, and sequenced using the Illumina NextSeq 500 platform using 75 cycles single end. This size range was selected to enrich for nucleosome-free regions.

### CRISPRa

CRISPRa was carried out as previously described ^6^. Briefly, gRNAs were designed in a region proximal to our new eTSS for *Mgat3* and prioritized based on off-targets/proximity to the eTSS. The *Mgat3* TSS (NCBI GeneID: 100689076) was detected using the annotation from ^34^ (https://www.synapse.org/#!Synapse:syn20999279.1). Target sequences and gRNA oligos are listed in Table S2 and Table S3, respectively. gRNAs were transfected along with a dCas9 fused with a VPR domain (VPR-dCas9) into mutant CHO-S cells carrying knockouts of *Mgat4a,4b* and *5, St3gal3,4* and *6, B3gnt2, Sppl3* and *Fut8* in biological triplicates. Non-targeting gRNAs were transfected with (NT-gRNA) and without VPR-dCas9 (NT-Cas9) as controls. Two days after transfection, cells were harvested to assess activation via qRT-PCR (in technical triplicate) and N-glycan analysis. Transcript levels were normalized to the mean of *Hprt* and *Gnb1* and relative expression levels were calculated using the 2^−ΔΔCt^ method ^35^. N-Glycans were fluorescently labeled and quantified via LC-MS.

### RNA processing

Sequence data for all RNA-seq data was quality controlled using FastQC (v0.11.6. Babraham Institute, 2010), and cutadapt v1.16 ^36^ was used to trim adapter sequences and low quality bases from the reads. Reads were aligned to the Chinese hamster genome assembly PICR ^9^ and annotation GCF_003668045.1, part of the NCBI Annotation Release 103. Sequence alignment was accomplished using the STAR v2.5.3a aligner ^37^ with default parameters. Reads mapped to multiple locations were removed from analysis.

### ATAC processing

Sequence data for ATAC-seq was processed using the ENCODE ATAC-Seq pipeline (https://github.com/kundajelab/atac_dnase_pipelines). The reads were trimmed using cutadapt v1.9.1. Reads were aligned using Bowtie2 v2.2.4 ^38^ to the same Chinese hamster genome. Peaks were called using Macs2 v2.1.0 ^39^ with a p-value of 0.01 and replicates were merged using irreproducible discovery rate (IDR) ^40^ of 0.1. The fold-change value is the number of normalized counts over the local background, taken as a 10,000 bp surrounding region.

### Peak Calling

#### HOMER peak calling

To call Transcription Start Site peaks, the Homer version 4.10 5’GRO-Seq pipeline was used (http://homer.ucsd.edu/homer/ngs/tss/index.html) ^27^. Briefly, aligned reads for TSS samples and control samples were estimated to have a fragment size of 1 base pair (bp). Counts, or tags, were normalized to a million mapped reads, or counts per million (CPM). Regions of the genome were then scanned at a width of 150 bps and local regions with the maximum density of tags are considered clusters. Once initial clusters are called, adjacent, less dense regions 2x the peak width nearby are excluded to eliminate ‘piggyback peaks’ feeding off of signal from nearby large peaks. Those tags are redistributed to further regions and new clusters may be formed in this way. This process of cluster finding and nearby region exclusion continues until all tags are assigned to specific clusters. For all clusters, a tag threshold is established to filter out clusters occurring by random chance. These are modelled as a Poisson distribution to identify the expected number of tags. An FDR of 0.001 is used for multiple hypothesis correction. Importantly, in experiments where the cap is enriched, efficiency is not perfect, and additional reads tend to occur in high-expressing genes. To correct for this, we use control samples, GRO-Seq and csRNA-input for GRO-Cap and csRNA-seq, respectively. These experiments do not enrich for the 5’ cap, and thus will be found along the gene body. We enforce our peaks to be more than 2-fold enriched compared to the controls. Motifs were visualized using HOMERs compareMotifs.pl ^41^.

### Processing Additional RNA molecules

GRO-cap and csRNA experiments can sequence RNAs beyond mRNA and lncRNAs, including unstable transcripts such as enhancer RNAs, spliceosomal snRNAs, and divergent transcripts which occur upstream and in the opposite direction of the coding mRNAs. Annotating these transcripts was outside the scope of our study and so we filtered them out by focusing on non-intronic RNAs and having a distance filter of 1 kb from the start of a transcript. However, these are kept in our release of all TSSs (not just protein-coding) for further use.

### Merging Samples

Sample peaks were merged using the mergePeaks command in Homer. Briefly, if samples have overlapping peaks, they are combined into one, where the start position is the minimum start position and the end is maximum end position. Additionally, when merging the samples’ peak expression in the same tissue, the average CPM was used.

### Promoter TSS calling

Since both TSS methods enrich for nascent transcripts, many peaks end up in enhancer regions, where enhancer RNA is transcribed. There were two annotations used in this study. The first is from the NCBI Annotation 103 release using the PICR genome. The other annotation ^9^ was used for designing the gRNAs in the CRISPRa experiment. That genome annotation used RNA-seq, proteomics and Ribo-seq to create a more complete annotation than NCBI. Therefore, to annotate protein-coding TSSs, a distance threshold from the original annotation was enforced. Ultimately, we used a distance of −1kb to +1kb from the initial reported TSS. Additionally, intron enhancers are known to occur, and so any intron peaks were filtered.

### RNA-Seq comparison

To compare RNA-seq to TSS-seq, we used 1,558 CHO samples of different lines that were a combination of in-house samples and public (see Table S4 for accession IDs). These were quantified and converted into transcript per kilobase gene per million mapped reads (TPM) using Salmon ^42^.

### Read histograms

For Figure 3A and 3B, Homer annotatePeaks.pl with the -hist command was used to construct the histogram. A bin size of 25 nucleotides was used, and the CPM per nucleotide per TSS was calculated. For example, with 1000 TSSs, 25 nucleotide bin size, and 2000 CPMs total within the bin, the height is 2000/(25 * 1000) = 0.08. Additionally, while it is expected to see many tags in one region, we restrict the number of tags to count per nucleotide to 3 to remove PCR duplicates.

### Motif Analysis

For motif analysis in Figure 2, the Homer command “findMotifsGenome.pl {bed file} {genome fasta file} {out_dir} -size -300,100 -len 6,8,10” was used.

Motif analysis of the core promoter elements the Initiator element and the TATA-Box seen in Figure 3 were done using FIMO of the MEME Suite 5.0.2 package with default parameters ^43^. Position weight matrices (PWMs) for the motifs were downloaded from Jaspar ^44^. To detect motifs, +/−1kb around each detected TSSs were extracted, and the PWMs were scanned across each position where a position score is the sum of the 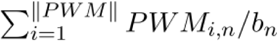 score, where i is the relative position in the genome and n is the nucleotide that corresponds to that position, and b is the background distribution for that nucleotide (assumed frequency of 0.25 for each nucleotide). These scores are summed and converted into a log_2_-likelihood ratio score for each motif with respect to each sequence position and then converts these scores to P-values, with a cutoff of 0.0001.

### Tissue-Specific Gene Enrichment Analysis (TSEA)

TSEA was done using the webserver http://genetics.wustl.edu/jdlab/tsea/. Log p-values of enrichment significance were used for Figure 2D.

### Peaks Naming and Ranking

Significant peaks identified were ranked by the total number of tags supporting each and then sequentially numbered (for example, p1@Gfap corresponds to the promoter of *Gfap* which has the highest support). This is similar to how the FANTOM Consortium reported their TSSs in mice ^45^. Additionally, if there is no peak for that gene, the original RefSeq location is used, and the name is numbered as 0 (for example p0@RRP7A). To rank peaks of the same transcript and gene, we accounted for both the number of samples and the expression value itself. For this, CPM values were first log transformed, and the median value of all samples with a peak was multiplied by the number of samples that had that peak:

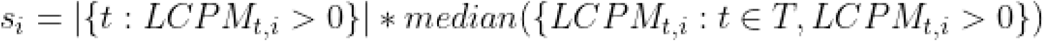

Where is the activation value for that peak in region *i, LCPM*_*t,i*_ is the log10 CPM in tissue *t* in TSS *i*, and T is the set of all tissues. The first term on the right-hand side is the number of biosamples that have a peak in this region, and the second term is the median value amongst those biosamples.

### Release of Annotations

A bed file with the TSSs will be released, with the ID as described above. This is done for both the NCBI RefSeq annotation GCF_003668045.1 and the synapse (https://www.synapse.org/#!Synapse:syn20999279.1) annotations. Each peak has an associated isoform and gene, and there may be more than one TSS per isoform. A companion metadata file, which is a tab-separated-value file where each row is a TSS corresponding to the TSS in the bed file. In the file the columns are as follow. (1) The name of the TSS that corresponds to the bed file IDs. (2) Comma-separated list of biosamples that express the TSS. (3) Confidence score with three binary values, where the leftmost bit is one if the maximum CPM is greater or equal to 2 CPM, the second bit is if there were more than one biosample with the TSS, and the third bit is 1 if it is considered a stable TSS, meaning there was total-RNA reads present there as well. (4) The location of the CHO ATAC region, if there is one near the TSS. (5) Corresponding gene. (6) Corresponding transcript. (7) If this TSS was annotated from the current paper or was from the previous annotation, (which is only there if there was no experimental TSS seen for that transcript). 8) Binary value if there was an open-chromatin region near the TSS in any of the samples, which includes more than just CHO (see Figure 1. for samples).

**Figure 1:**
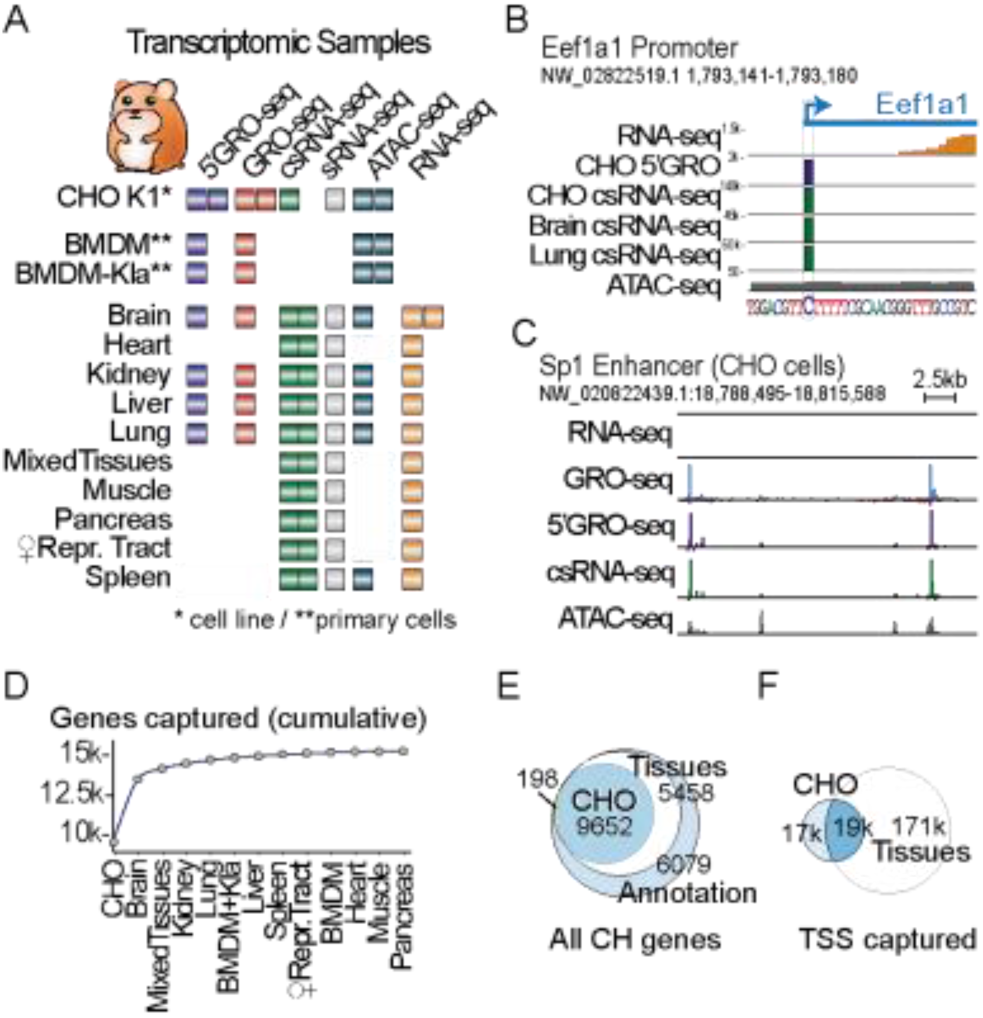
A Chinese hamster Transcriptome Atlas. (A) Overview of datasets generated here. (B) Example transcription start site at single-nucleotide resolution as defined by 5’GRO-seq and csRNA-seq (using GRO-seq and sRNA-seq as input, respectively) of the focused Eukaryotic Translation Elongation Factor 1 Alpha (Eef1A1) promoter in CHO cells and diverse tissues. RNA-seq reads are shown in orange. (C) Example of unstable transcription start sites of enhancer RNAs that are poorly detected by conventional RNA-seq at the Sp1 “super enhancer” locus in CHO cells. Note: Raw IGV browser visualization data are provided in Supplementary Figure S3. (D) Cumulative plot of all mRNA genes TSSs across all samples as detected by csRNA-seq and/or 5’GRO-seq enrichment over GRO-seq and/or csRNA-seq. Sorted by taking CHO as the first sample, then include the next sample to add the most additional genes. (E) Detection of TSSs (CPM <1) in CHO and hamster samples of all NCBI annotated genes. 198 is the overlap between CHO and the annotation, excluding the hamster tissues. (F) Venn Diagram of the total TSSs captured by 5’GRO and csRNA-seq in CHO K1 cells and the tissues using an FDR of 0.001, including putative non-coding RNA TSSs and enhancer RNA TSSs.

**Figure 2:**
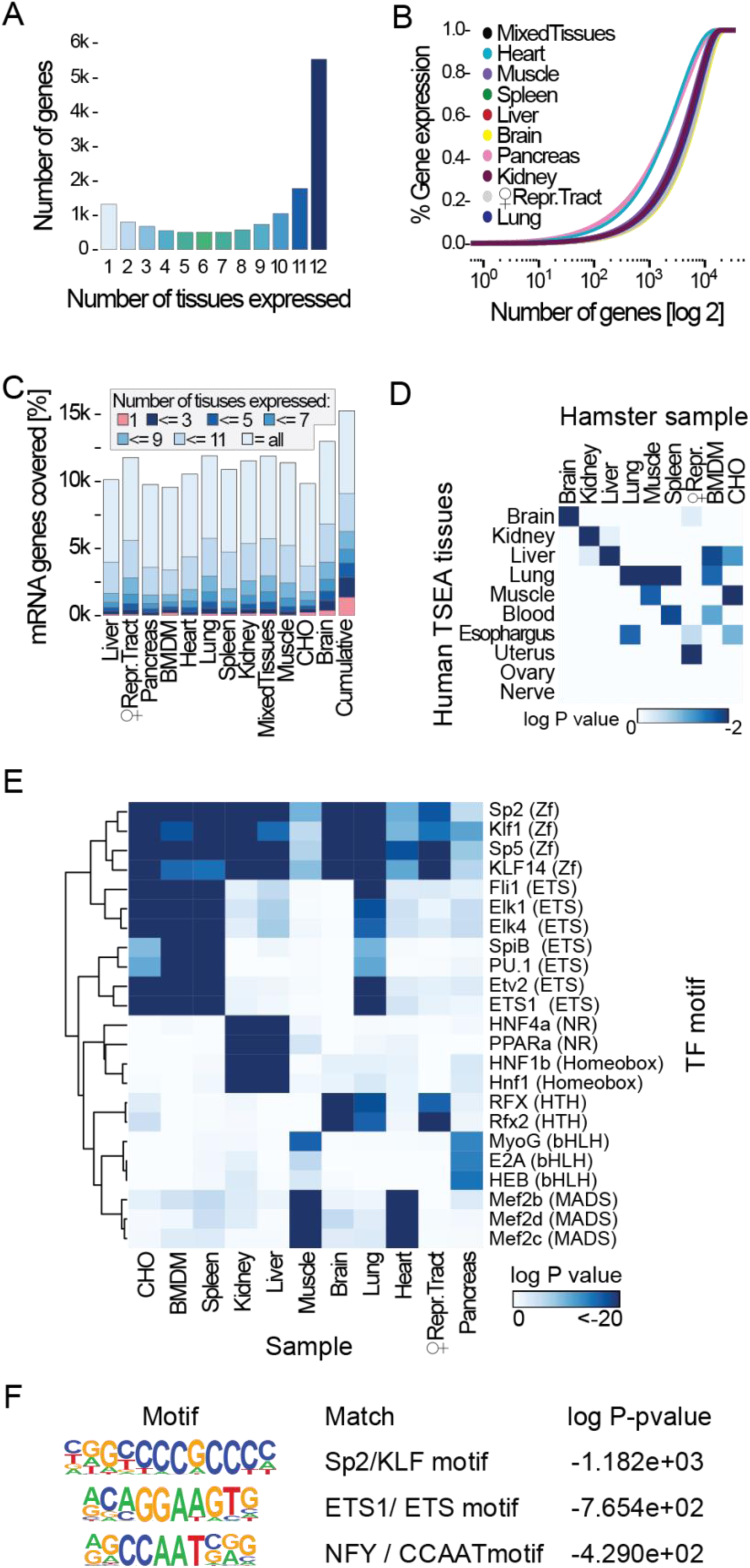
Composition and complexity of diverse tissue-specific Chinese hamster transcriptomes. (A) Experimentally detected genes and the number of tissues they were confidently expressed as defined by csRNA-seq and 5’GRO-seq. (B) Cumulative plot of the distribution of transcript abundance as defined by RNA-seq in various tissues. The transcriptome of highly specialized tissues such as the heart, muscle or the pancreas is more dominated by the high expression of a small set of specific RNAs than those of complex tissues such as the brain. (C) Comparison of gene expression distributions across tissues as defined by csRNA-seq and 5’GRO-seq. (D) Tissue-specific gene enrichment analysis (TSEA) comparing the gene expression patterns of our samples as defined by csRNA-seq and 5’GRO-seq to orthologous human pre-defined tissue-specific genes. (E) Transcription factor motifs (top 3 per sample) enriched across the tissue-enriched TSSs in each tissue highlight conservation and factors involved in maintaining tissue-specific expression patterns. Enrichment defined as a TSS in a tissue that has CPM 3 standard deviations above the mean of each respective TSS. (F) Transcription factor motifs enriched in all TSSs in the eTSS annotation.

**Figure 3:**
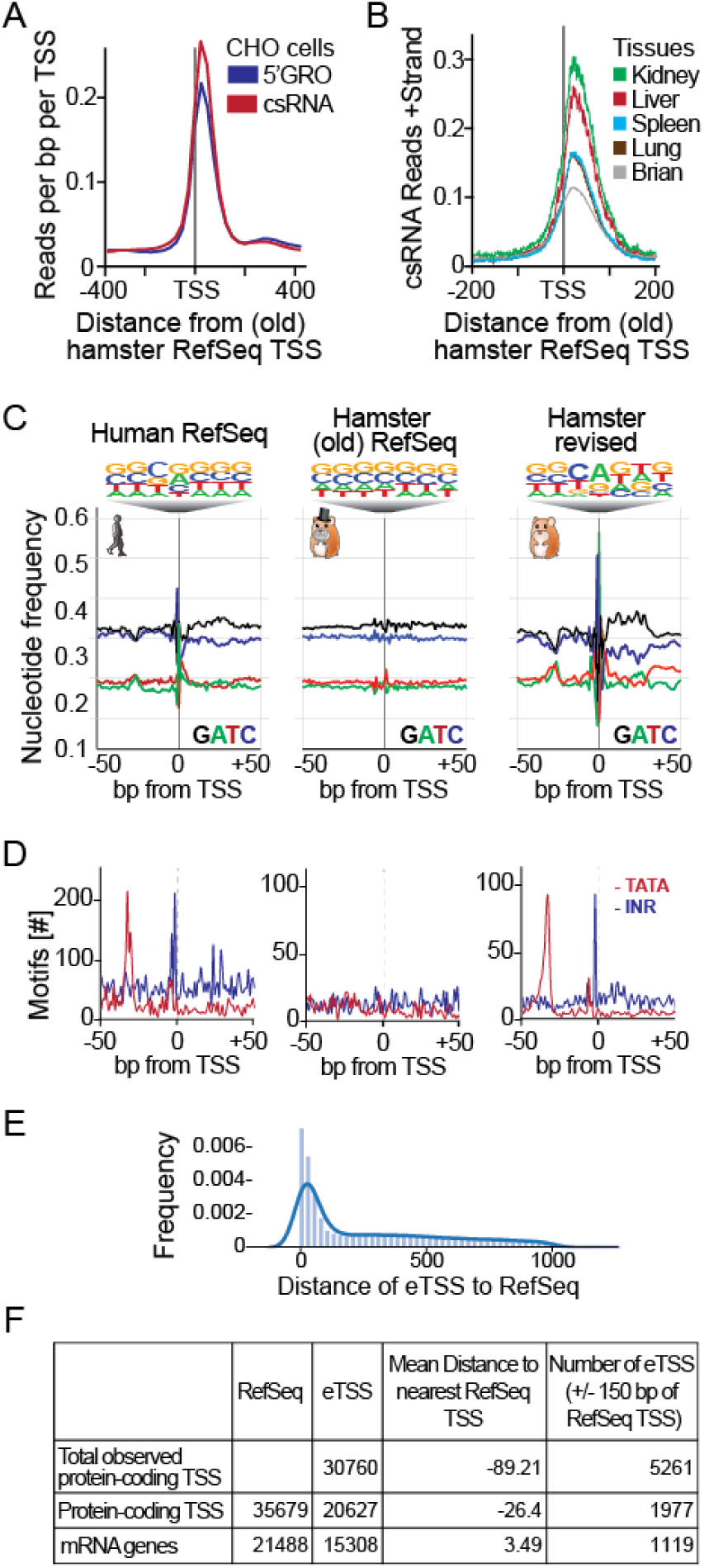
An experimental realignment of TSS annotation for the Chinese hamster uncovers expected genomic elements. (A) Comparison of experimentally defined TSSs from CHO cells by 5’GRO-seq and csRNA-seq relative to the current Chinese hamster (GCF_003668045.1) annotation. (B) Comparison of experimentally defined TSSs from selected tissues relative to the current Chinese hamster (GCF_003668045.1) annotation. (C) Nucleotide frequency plots of TSSs and their relative information content in Human RefSeq, Chinese hamster (GCF_003668045.1), and our revised Chinese hamster annotation. (D) Frequency distribution of positional core promoter elements: the TATA box and the Initiator that are commonly found at −30 and +1, relative to the TSS. (E) Distribution of distance between eTSSs observed and the nearest NCBI RefSeq TSS. (F) Statistics of total protein-coding TSSs observed and their distance to NCBI RefSeq TSSs. For rows 2 and 3, distance is the nearest distance of an eTSS to an NCBI transcript, taking the minimum in that NCBI transcript or gene, respectively.

### Peak Location

When samples are merged together, peaks may be offset by a few base-pairs, either due to noise or having multiple peaks within the 150 base-pair window used to call peaks. Although all of these are near each other, we report here the location from only one sample to avoid nucleotide shifting. When reporting the final location we use one of two options: if the peak was seen in CHO, we report the CHO location, otherwise, we report the location of the peak in the sample which had the largest CPM.

### Open Chromatin

In addition to peak location, the presence of the TSS within 250 bps of open chromatin is reported for both the CHO cell line and in general across all samples tested. The former may be of use when trying to engineer CHO, and the latter adds credibility to the TSS found.

## RESULTS

### Nascent and steady state RNA sequencing yields nucleotide-resolution of diverse RNA classes and TSSs across tissues

Algorithms predicting gene annotations rely on highly conserved features such as protein domains. Consequently, although genes are commonly assigned correctly, their TSSs and promoter annotations are often inaccurate, as these features evolve rapidly and can relocate to non-homologous regions ^46^. To correctly annotate TSSs in protein genes and non-coding RNAs (e.g. pri-miRNAs, lncRNAs, snoRNAs), it is necessary to experimentally determine these features. Thus, we cultured CHO-K1 cells and acquired female hamsters from Dr. George Yerganian, representing the original colony from which CHO cells were derived in 1957 ^47^. We comprehensively profiled their coding and non-coding transcriptomes using multiple nascent and steady-state assays (Figure 1B, Supplementary Figure S1A); assays were conducted on ten tissues and bone marrow derived macrophages (BMDMs) in resting state and after 1h stimulation with Kdo2-Lipid A (KLA; activates BMDMs by mimicking a bacterial infection ^48^). This provided a comprehensive view of diverse RNA classes and genomic elements, including for genes typically silenced in CHO cells.

Accurate TSSs are best determined by nascent RNA sequencing methods to avoid capturing potential false-positives 5’ends caused by RNA processing or recapping of cytosolic (steady-state) mRNAs ^49,50^. It is also preferred to adopt methods that selectively focus the sequencing power to the 5’end of transcripts. Even for highly expressed genes, such as the Eukaryotic Translation Elongation Factor 1 Alpha (Eef1a1, Figure 1B), RNA-seq and other methods that capture the complete transcriptome have limited information about the exact location where genes start and often fall short in the detection of the TSSs for less abundant transcripts. Thus, when feasible, we captured nascent transcription using global nuclear run on sequencing (GRO-seq) ^25^ and nascent TSSs by 5’GRO-seq ^26^ (also known as GRO-cap ^50^).

However, as GRO-seq requires the purification of several million cellular nuclei, which is not always readily feasible for tissues or preserved samples, we also took advantage of capped small RNA-seq (csRNA-seq). csRNA-seq accurately captures initiating transcripts and thus the TSSs largely independent of transcript stability, similar to 5’GRO-seq, but uses total RNA with RINs as low as 1 as input ^27^. The flexible input material requirements and comparatively simple csRNA-seq protocol enabled us to expand our analysis across diverse hamster tissues.

TSSs leading to the production of stable RNAs are a hallmark of genes and non-coding RNAs. TSSs producing unstable or transient RNAs associate with pri-miRNAs and enhancer regions (Figure 1C) ^51,52^. The TSSs captured among 5’GRO-seq replicates and between 5’GRO-seq and csRNA-seq were highly consistent in their position and expression strength (Pearson correlation of r=0.96 and r=0.88, respectively, Supplementary Figure S2). Identification of these high quality TSSs were achieved in part by employing our GRO-seq, and small RNA-seq (sRNA-seq) data as a background control (also known as input) for 5’GRO-seq and csRNA-seq, respectively (Supplementary Figure S1B). To confirm our TSSs and confidently distinguish stable and unstable TSSs, we integrated areas of accessible chromatin from ATAC-seq ^28^ and measured steady-state gene expression by conventional ribosomal RNA-depleted RNA-seq (Supplementary Figure S1B). A complete list of the 73 datasets generated and used in our transcriptome analysis is provided in Supplementary Table S1. Together, these data provide accurate TSSs at single-nucleotide resolution and a comprehensive view of that nascent and steady-state hamster transcriptome (Figure 1, Supplementary Figure S3 A-C).

### Profiling diverse hamster tissues identifies TSSs for silenced genes in CHO cells

While CHO cells are exceptional protein production hosts, many genes that could improve product quality or quantity lay dormant. Indeed, about 50% of genes, including many that contribute to important human post-translational modifications, are silent ^21^. Consistent with previous findings, we detected TSSs for only 46% of protein-coding genes using nascent TSS mapping in CHO cells (Figure 1E). By integrating our TSSs from ten tissues and resting and stimulated BMDMs, we confidently recovered TSSs from an additional 5,464 protein-coding genes (Figure 1E) (13,666 overall protein-coding TSSs), 2,657 non-coding RNAs, 63 miRNAs and 154,736 putative distal regulatory elements (enhancer RNAs), totaling to over 177,000 new TSSs (Figure 1C-F). Together, these data provide experimentally determined TSSs for 70% of the protein-coding genes. Functional gene groups that were less covered by our data include those associated with olfaction and taste, the male sex organ (testis), development and the adaptive immune system (Supplementary Figure S3 D,E). In addition, we identified alternative TSSs for 55% of the proposed alternative protein isoform start sites (Supplementary Figure S4). This isoform annotation is important as it facilitates the tailored expression of protein isoforms that can exhibit differential activity or distinct functions ^53,54^.

### Tissue-specific TSS and gene expression patterns in the Chinese hamster

Capturing the active (nascent) and steady-state transcriptome across tissues and cell lines revealed that about 1/3 of annotated genes were ubiquitously expressed, while only a comparatively small number of genes were tissue-specific (Figure 2A). The apparent transcriptome complexity ranged from 9,596-9,850 detected genes in BMDMs (pooled rested and stimulated conditions) and CHO cells, respectively, to 9,782 genes confidently detected in the pancreas and 13,007 in the brain (FDR=0.01, see Material and Methods and Figure 2B). This observed complexity is related to the tissues’ degree of specialization and the number of different cell types found within the tissue, but also affected by high abundance transcripts that can hinder detection of less abundant ones ^55^. In the pancreas, for example, much of transcription is directed towards expressing secretory enzymes such as chymotrypsinogen or carboxypeptidase ^56^, while in the brain, a higher diversity of transcripts are expressed^57,58^.

Of the genes for which transcription was confidently detected (n = 15,308; ∼71% of all genes annotated in RefSeq), 40% were shared among all 12 tissues or cell types and another 19% were found in more than 11 samples. Approximately 8% of genes were unique to one tissue or cell type which increased to 18% and 25% for genes expressed in less than 3 or less than 5 samples, respectively (Figure 2C). These tissue-specific gene expression signatures were enriched for genes previously annotated as unique in the analogous human tissues, as determined by Tissue-Specific Gene Enrichment Analysis (TSEA,^59^, Figure 2D). Thus, although thousands of genes are differentially expressed between tissues, a limited number of genes were exclusive to a tissue or cell type (Figure 2A,C).

To gain insights into the regulatory programs underlying the diversity of the analyzed samples, we next investigated the promoters of the expressed genes. Notably, although most genes were expressed across several tissues and cell types (Figure 2A), the promoters and initiation sites within these promoter regions often varied, leading to remarkable diversity in 5’ ends for most genes (S4B-C). This finding highlights regulatory plasticity as a critical factor to maintain gene expression as regulatory programs diverge to support distinct cell types. To explore which transcription factors may drive these tissue-specific gene expression programs in the Chinese hamster, we next probed tissue-specific gene promoters for differentially enriched transcription factor binding motifs. This analysis highlighted known key regulators or lineage determining transcription factors with preferential expression and binding sites for each tissue such as Rfx factors for the brain ^60^, HNF1 ^61^ and PPARα factors for the kidney and liver ^62,63^ or the MADS-box transcription factors Mef2b,c and d for the heart and muscle ^64,65^. The top 3 enriched motifs from each sample are shown in Figure 2E. Closely related tissues such as muscle and heart or liver and kidney displayed a combination of shared and unique factors, which also became apparent for other tissues once more motifs were integrated into the analysis. This observation is in line with the hypothesis that tissue-specific regulatory pathways arise by tinkering with existing pathways, rather than complete innovation ^66,67^ of regulatory elements needed. On the other hand, ubiquitously expressed genes were enriched for the binding motifs of strong, ubiquitous activators such as SP2/KLF family members ^68^, ETS factors or NFY (Figure 2F). Together, these findings show that a comparatively large fraction of genes, including ubiquitous transcription factors, ensure the cell’s vital core programs, while a comparatively small number of genes can effectively facilitate specialization. Moreover, our identification of tissue-specific genes and transcription factors enriched in their promoters are consistent with other mammals, further validating our experimental TSSs and RNA-seq data.

### Realignment of Chinese hamster TSSs exposes key features of the core promoter

Genome annotations are an essential part of many sequencing and bioinformatic analyses. We therefore integrated our experimental data to refine and revise the annotations for the Chinese hamster TSSs. This is referred to as experimental TSS annotation (eTSS). First, we identified high confidence single-nucleotide TSSs (Supplementary Figure S1B) that were either consistent across both 5’GRO and csRNA-seq, or across several tissues or samples (see Material and Methods for details). These TSSs were also consistent with open chromatin regions, as defined by ATAC-seq, and typical histone modifications for active promoters (Supplementary Figure S5). To distinguish stable and unstable transcripts, we integrated the previous gene annotations with our RNA-seq data from tissues. Stable transcripts were considered if there were RNA-seq reads within 150 bps. Comparison of our annotation with other RNA-seq datasets enables the assessment of false positive and false negative TSSs in our revised annotation. Thus, we analyzed 1,558 CHO RNA-seq samples ^21,69–71^. Genes where we failed to experimentally detect a TSS showed little to no expression across the CHO RNA-seq datasets while those where we captured a TSS were consistently expressed (Supplementary Figure S6).

We next evaluated the relationship of our experimental TSSs to the Chinese hamster RefSeq TSSs (GCF_003668045.1). Many of the RefSeq annotations were computationally predicted with support of RNA-seq but not TSS data. TSSs independently mapped by both 5’GRO-seq or csRNA-seq displayed a similar distribution (Figure 3A). However, the experimentally determined TSSs displayed a clear offset from the RefSeq annotation. A comparable offset was also observed for TSSs measured in diverse tissues (Figure 3B).

To more independently assess the differences between RefSeq and our revised annotation, we plotted their TSS proximate nucleotide frequency distributions. Basal transcription factors often bind core promoter elements to recruit and position the RNAP II transcription complex which preferentially initiates on purines ^72–74^. These nucleotide preferences are clearly visible when analyzing the human RefSeq (GRCh38) annotation and in our revised hamster annotation, but not in the current Chinese hamster RefSeq annotation (Figure 3C). In addition to the increased information content in the TSS-proximate nucleotide frequencies, the TATA box and Initiator (Inr) core promoter elements ^75–77^, were found at the expected −30 and +1 bp positions respectively in the human RefSeq and in our revised Hamster annotations, but not the old RefSeq annotation (Figure 3D). Overall, most eTSSs differed from the NCBI RefSeq annotation, and on average, an eTSS is 89 nucleotides upstream from an NCBI TSS (Figure 3E, 3F). Together, these data provide an independent validation for our improved annotation and stress the importance of experimental TSS data for accurate genome annotations.

### Accurate TSSs facilitate the activation of dormant genes for engineering of CHO gene expression

CRISPR/Cas9 enables rapid and cost-effective genome editing, gene inhibition (CRISPRi), and activation (CRISPRa) without altering the native DNA sequence ^19,20,22^. However, the success of these and similar precise genome engineering approaches ^18,78^ depends on accurate gene annotations. To demonstrate the utility of our refined eTSS annotation for the Chinese hamster, we investigated genes associated with glycosylation, given the majority of protein therapeutics are glycosylated, and the glycans can significantly impact drug safety, efficacy, and half-life ^79^. We detected dozens of eTSSs across diverse classes of glycosylation enzyme genes (Figure 4A). We focused specifically on *Mgat3* (Figure 4B), which is naturally dormant in CHO cells ^80^, leading to a lack bisecting N-acetylglucosamines on glycoproteins, as seen on some human glycoproteins ^6,81^. Bisecting N-acetylglucosamines are important role in regulating complex glycosylation maturation and impact antibody effector function ^82,83^. Furthermore, *Mgat3* has previously been targeted by CRISPRa using the CHO RefSeq TSS ^6^, thereby providing an attractive gene for direct comparison.

**Figure 4:**
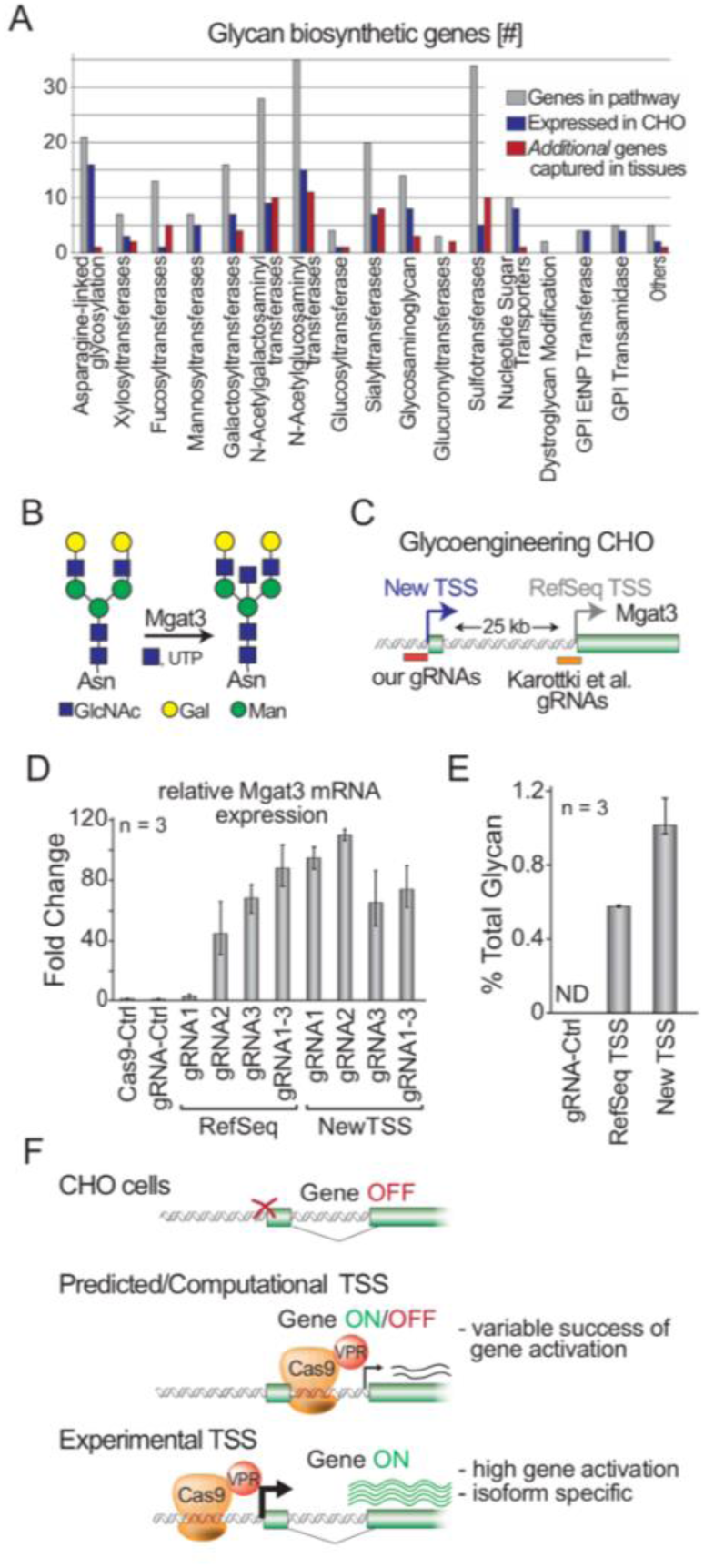
Experimentally-measured TSSs facilitates precision genome engineering to humanize glycosylation. (A) List of classes of glycosyltransferases in the Chinese hamster with the number of genes expressed in CHO cells (blue) and additional genes for which experimental TSSs were discovered in our tissue samples. (B) The *Mgat3*-encoded glycosyltransferase catalyzes the addition of bisecting N-acetylglucosamines on glycoproteins, but is silenced in CHO cells. (C) Overview of the experimental, new TSS and the RefSeq TSSs targeted by CRISPRa to induce *Mgat3* expression in CHO cells. (D) Quantitative RT-PCR measurement of *Mgat3* expression in CHO cells and upon activation by the three designed gRNAs using either NCBI RefSeq’s TSS or our new eTSS. As a control, the cells were transfected with NT-gRNA (gRNA-Ctrl) or NT-gRNA and VPR-dCas9 (Cas9-Ctrl) (E) Comparison of the levels of bisecting N- acetylglucosamines following CRISPRa. As a control, the cells were transfected with NT-gRNA (gRNA-Ctrl). (F) Overview: Experimental TSSs facilitate efficient engineering gene expression pathways.

To activate *Mgat3* we designed three CRISPR guide RNAs (gRNAs) complementary to the DNA sequence starting at −641, −522 and +6 bps with respect to our experimental TSS, (gRNA 1, gRNA 2, & gRNA 3 respectively), which was approximately 25 kb away from the NCBI TSS (Figure 4C). CRISPRa resulted in a mean of 94-,109-, and 64-fold upregulation for gRNAs 1, 2, and 3, respectively (n=3 samples each) of Mgat3 as measured by qRT-PCR, and 73-fold for a mixture of the 3 (Figure 4D). Despite being 25,481 bp upstream, targeting this new TSS resulted in activation that was comparable to that observed when previously targeting the RefSeq TSS^6^, To test if the resulting *Mgat3* transcripts impact glycan synthesis, we measured the relative % of glycans on the secretome that contained N-linked glycans with bisecting N-acetylglucosamine after targeting *Mgat3* at the RefSeq TSS and the eTSS. For this comparison we utilized the gRNAs that resulted in the highest activation levels – 109 and 88 fold for eTSS and RefSeq TSS, respectively. Activation with the RefSeq TSS resulted in an average of 0.58% of the total glycans containing the bisecting N-acetylglucosamine while the eTSS averaged 1.08% (Figure 4E). Together, these data show that our eTSS, which localized approximately 25 kb upstream from the predicted TSS, successfully activated and produced a functional *Mgat3*-encoded enzyme. With our newly reported TSSs for 15,308 genes, we anticipate further usage of these TSSs for gene engineering.

## DISCUSSION

In this study we measured and analyzed the coding and non-coding RNA in the Chinese hamster genome using steady state and nascent RNA sequencing experiments for diverse hamster tissues and cell lines. Through this we were able to comprehensively map TSSs for >70% of annotated Chinese hamster genes and non-coding RNAs, including many genes normally silenced in CHO cells. Furthermore, we identified tissue-specific TSSs, including many for genes silenced in CHO cells. Importantly, we have been able to realign current TSSs in RefSeq, which were predominantly computationally predicted. The experimentally-measured TSSs uncovered expected features of the core promoter and we demonstrated it can be used to activate a silenced gene of interest using CRISPRa. Through this we present an invaluable resource to guide genome editing and genomic analysis of CHO cells.

Here we captured 25,442 nascent protein-coding TSSs (CPM >1) corresponding to 15,308 genes, along with 4,478 ncRNAs (lncRNA, miRNA, snRNA, snoRNA, and tRNA), and 176,914 distal peaks. This resource provides rich information for precise gene engineering. Furthermore, including diverse hamster tissues will help in efforts to fine tune existing CHO gene regulatory programs, and activate genes or pathways naturally encoded in the Chinese hamster genome but dormant in CHO cells. Our TSSs are a prerequisite for the design and testing of gRNAs and eventually, an effective gRNA library for the activation of diverse Chinese hamster genes by CRISPRa (Figure 4F). It can also complement existing data on epigenetic markers of CHO cells in efforts to find endogenous promoters that avoid silencing seen with common viral promoters or harness endogenous regulatory circuits involved in ER stress or cold shock ^84^.

Our transcriptomic datasets also provide a comprehensive resource for future research and discovery. In addition to our gene-centric atlas of Chinese hamster TSSs reported here, our data cover a plethora of transcriptomic features as well as other transcripts such as pri-miRNAs or those originating from distal regulatory elements that are commonly referred to as enhancer RNAs. Although beyond the focus of this manuscript, this extensive, transcript stability-independent resource of TSSs should aid to improve our understanding of how gene expression is regulated in hamsters and how tissue-specific regulatory programs emerged. While a key advantage of CRISPRa is the ability to activate desired genes independent of tissue-specific transcription factors, future engineering efforts may be more tailored towards adjusting transcriptional programs, rather than one or a few specific genes. For example, our definition of transcription factors that were highly enriched in the promoters of tissue-specific genes provides a first step to advance our understanding of which and why specific genes or pathways are silent in CHO cells. Improved knowledge of how gene regulatory networks function in hamsters may ultimately allow to predict how activation of one gene impacts the hamster regulome and to eventually finetune desired regulatory programs, rather than individual genes ^85^. Going beyond capturing TSSs, our data also contain maps of open chromatin, as defined by ATAC-seq, nascent transcription, as defined by GRO-seq, and mature RNAs, as defined by ribosomal RNA-depleted RNA-seq for CHO cells, hamster macrophages and diverse hamster tissues which were primarily used in this study as a critical input for the identification of high-confidence TSSs. Our data thus also provide a rich resource for future studies and enable the integration of the Chinese hamster into comparative or evolutionary studies, for example, as an outgroup to mice ^31^.

In summary, our data has enabled the development of a compendium of experimentally defined TSSs and transcriptomic features from multiple tissues and cells types from the same hamster colony from which CHO cells were generated. Our revised annotation shows considerable improvement over the current RefSeq by several measures including agreement with published RNA-seq datasets, TSS information content as well as core promoter motifs. More broadly, these findings emphasize the importance of refined TSS mapping methods such as 5’GRO-seq/GROcap or csRNA-seq for accurate annotation of a gene’s 5’ end. The TSS is a landmark in gene regulation and its accuracy becomes imperative in an era of genetic engineering. We further envision that our data and annotation will provide a rich resource for the CHO community and beyond as the Chinese hamster is further included in comparative and evolutionary studies. At its core though, the improved TSSs map will aid CHO gene engineering efforts aiming to improve the quality and quantity of desired recombinant proteins and ultimately reduce drug manufacturing costs.

### DATA AVAILABILITY

The transcription start site annotation, is released as updates to both NCBI’s RefSeq annotation and Synapse annotation (synapse ID [syn20999279.1](https://www.synapse.org/#!Synapse:syn20999279)), and each comes with a bed file containing the location of the TSSs, along with two accompanying tab-separated metadata files. The genome used was *Cricetulus Griseus* genome version PICR GCF_003668045.1. All sequencing data are submitted to the Gene Expression Omnibus (GEO) with GEO ID GSE159044. Additional processed files are listed in Table S1, which is the GEO submission metadata sheet. The annotation files are also uploaded to Synapse (synapse.org), with ID syn22969187.

## SUPPLEMENTARY DATA

Supplementary Data are available at NAR online.

## ACKNOWLEDGEMENT

Marten A. Hoeksema for culturing BMDMs

## DEDICATION

The authors dedicate this work to Dr. George Yerganian (1924-2019), who provided the hamsters for this study, and for the original CHO cells in 1957.

