## Supplementary Figures for "A Chinese hamster transcription start site atlas that enables targeted editing of CHO cells"

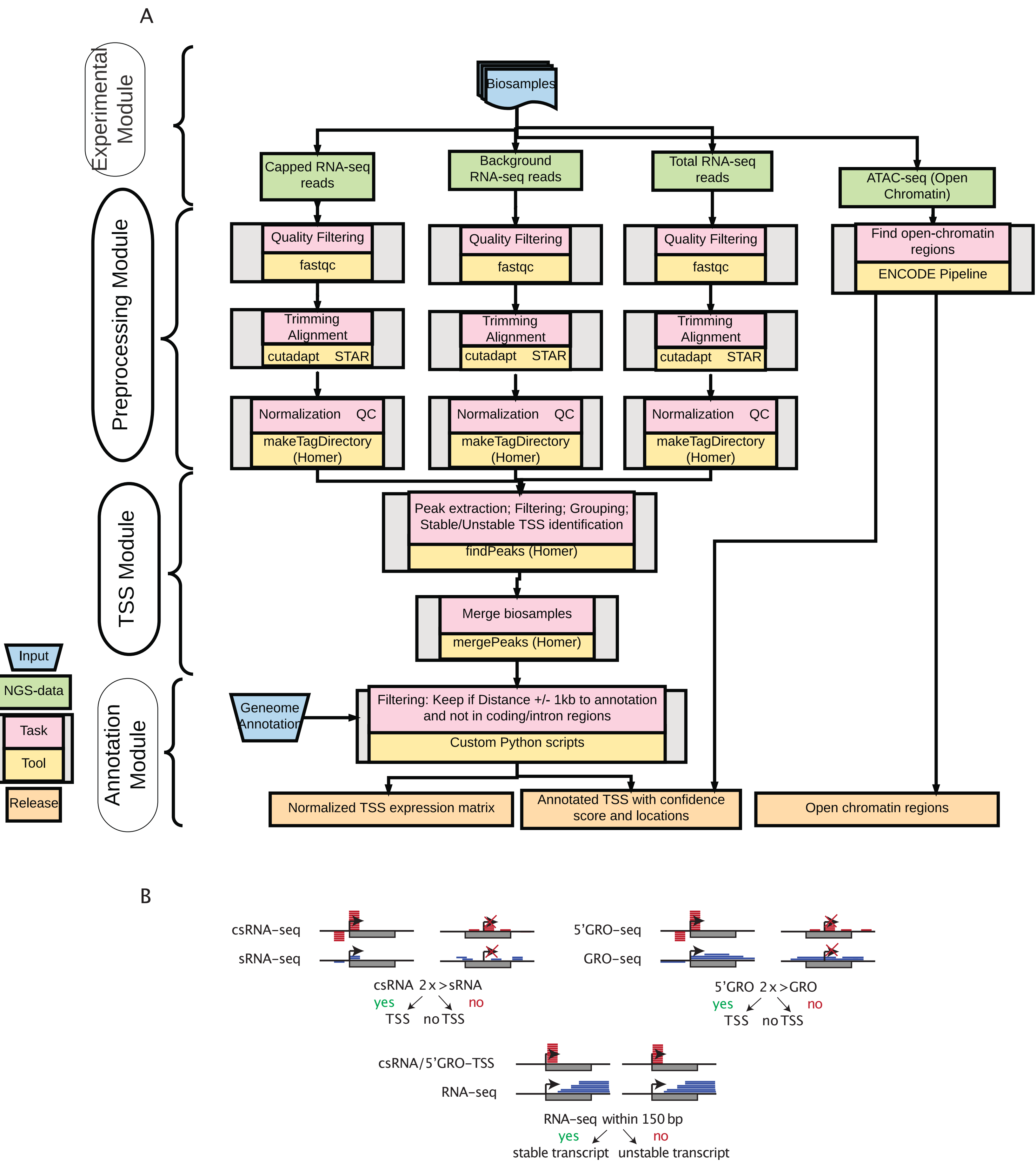


Figure S1
(A) Bioinformatics pipeline for TSS annotation. This shows the steps required along with the commands used to run these steps. Background RNA–seq reads are the sRNA–seq and GRO–Seq. Total RNA–seq reads are used for assessing if transcripts are stable or unstable, which is added in the confidence scores.(B) Scheme of transcription start site identification for 5'GRO–seq and csRNA–seq.

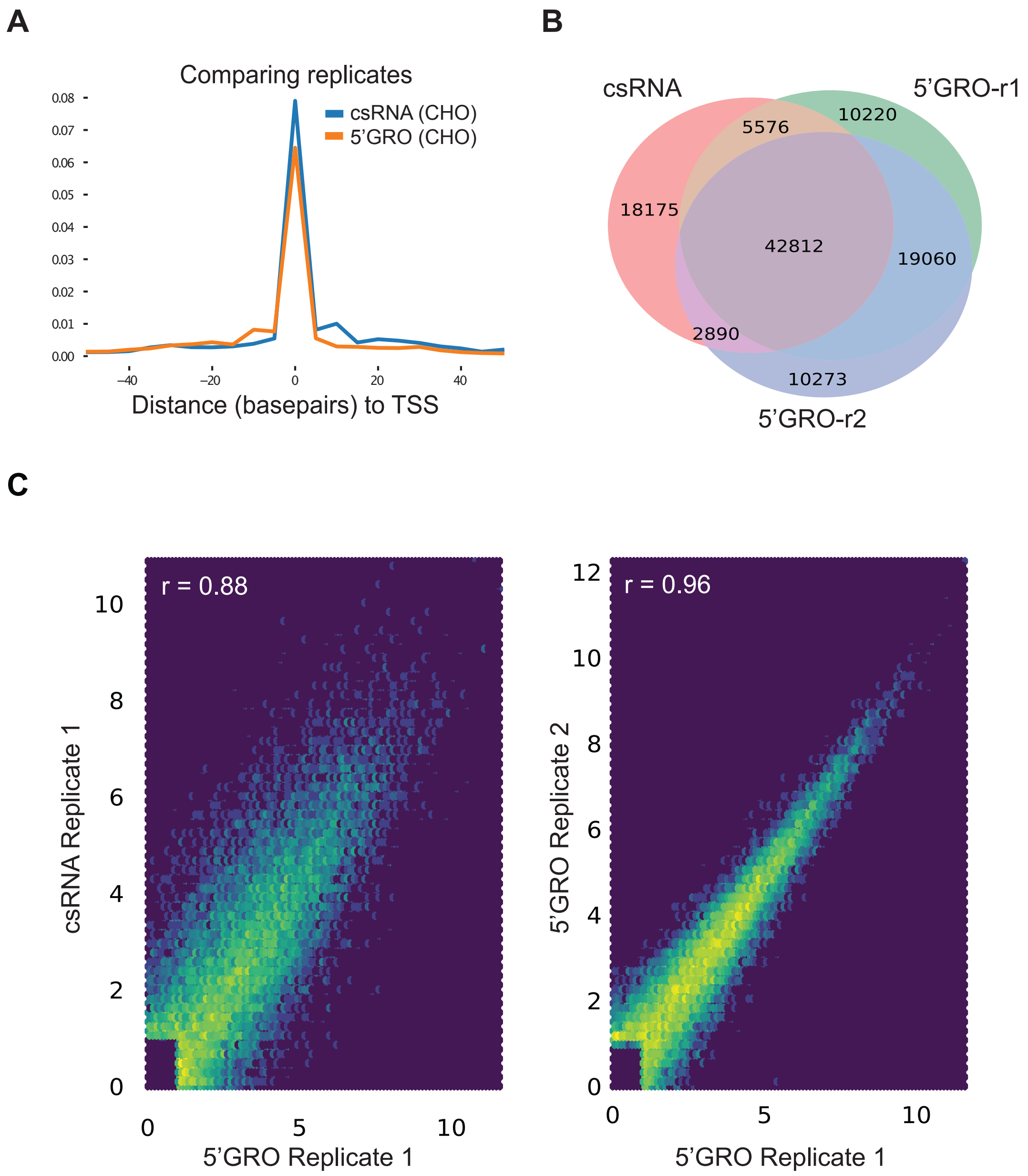


**Figure S2:**
(A) Density plot of the distance between peaks in different CHO replicates relative to a CHO GROcap sample B. CHO csRNA and GROcap sample A are shown nearby. (B) Number of overlapping total TSSs across all CHO replicates. (C) Density scatterplot of CHO replicates. Values are in log2 CPM. Left: CHO GRO-cap A vs csRNA-seq (pearson r=0.88, p-value < 0.001) Right: CHO GRO-cap replicates. (r=0.96, p-value < 0.001).

A

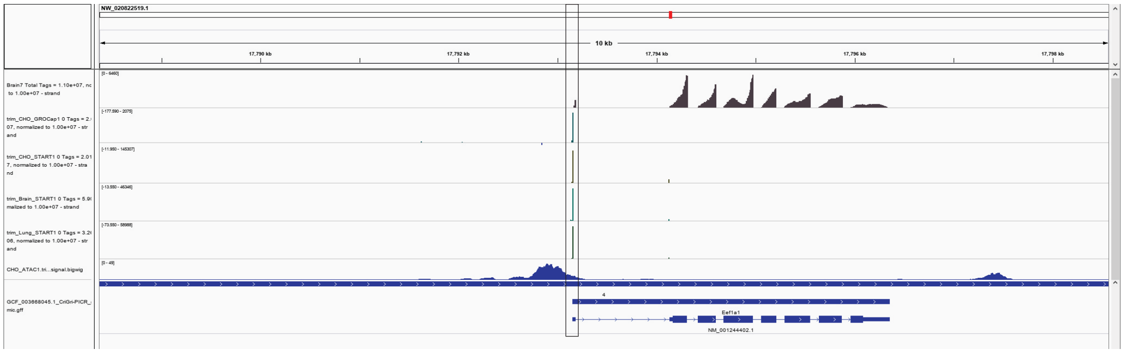

B

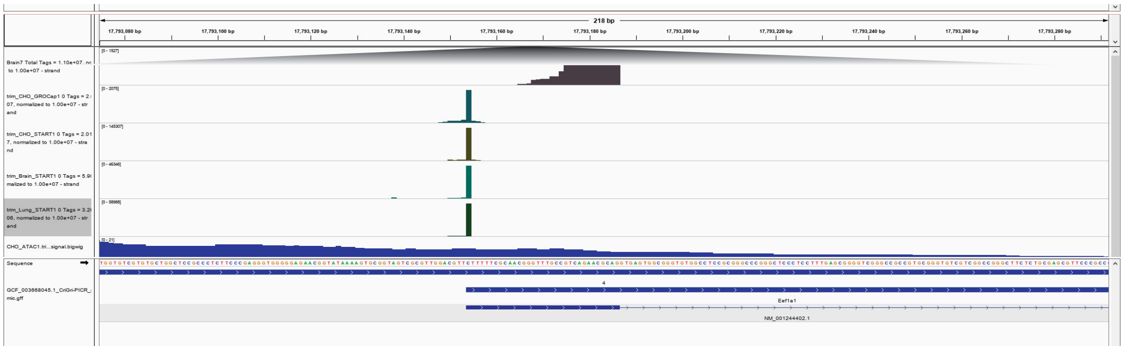

C

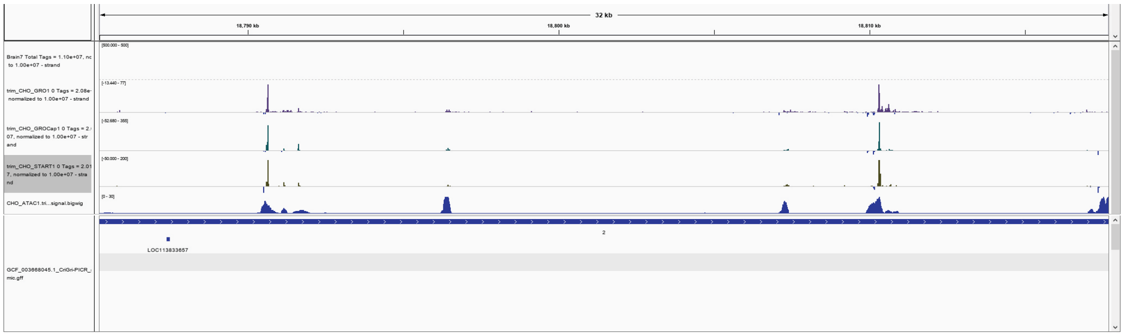

D

| Sublist | Category | Term | RT | Genes | Count | % | P-Value | Benjamini |
|---|---|---|---|---|---|---|---|---|
| | KEGG_PATHWAY | Olfactory transduction | RT | | 1040 | 22.1 | 0.0E0 | 0.0E0 |
| | KEGG_PATHWAY | Taste transduction | RT | | 31 | 0.7 | 2.3E-6 | 3.2E-4 |
| | KEGG_PATHWAY | Autoimmune thyroid disease | RT | | 39 | 0.8 | 1.3E-3 | 1.1E-1 |
| | KEGG_PATHWAY | Neuroactive ligand-receptor interaction | RT | | 103 | 2.2 | 1.5E-3 | 1.0E-1 |
| | KEGG_PATHWAY | Jak-STAT signaling pathway | RT | | 53 | 1.1 | 3.7E-2 | 8.8E-1 |
| | KEGG_PATHWAY | Cytokine-cytokine receptor interaction | RT | | 68 | 1.4 | 4.8E-2 | 9.0E-1 |
| | KEGG_PATHWAY | Ribosome | RT | | 73 | 1.5 | 7.7E-2 | 9.6E-1 |

E

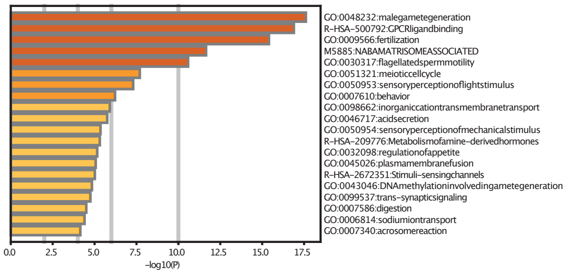


**Figure S3**
(A) IGV browser shots from Eukaryotic Translation Elongation Factor 1 Alpha 1Gene and (B) promoter. (C) IGV browser shot of the Sp1 “super enhancer”. (D) Enriched pathways and gene ontology (E) of genes for which TSSs were not detected in our samples.

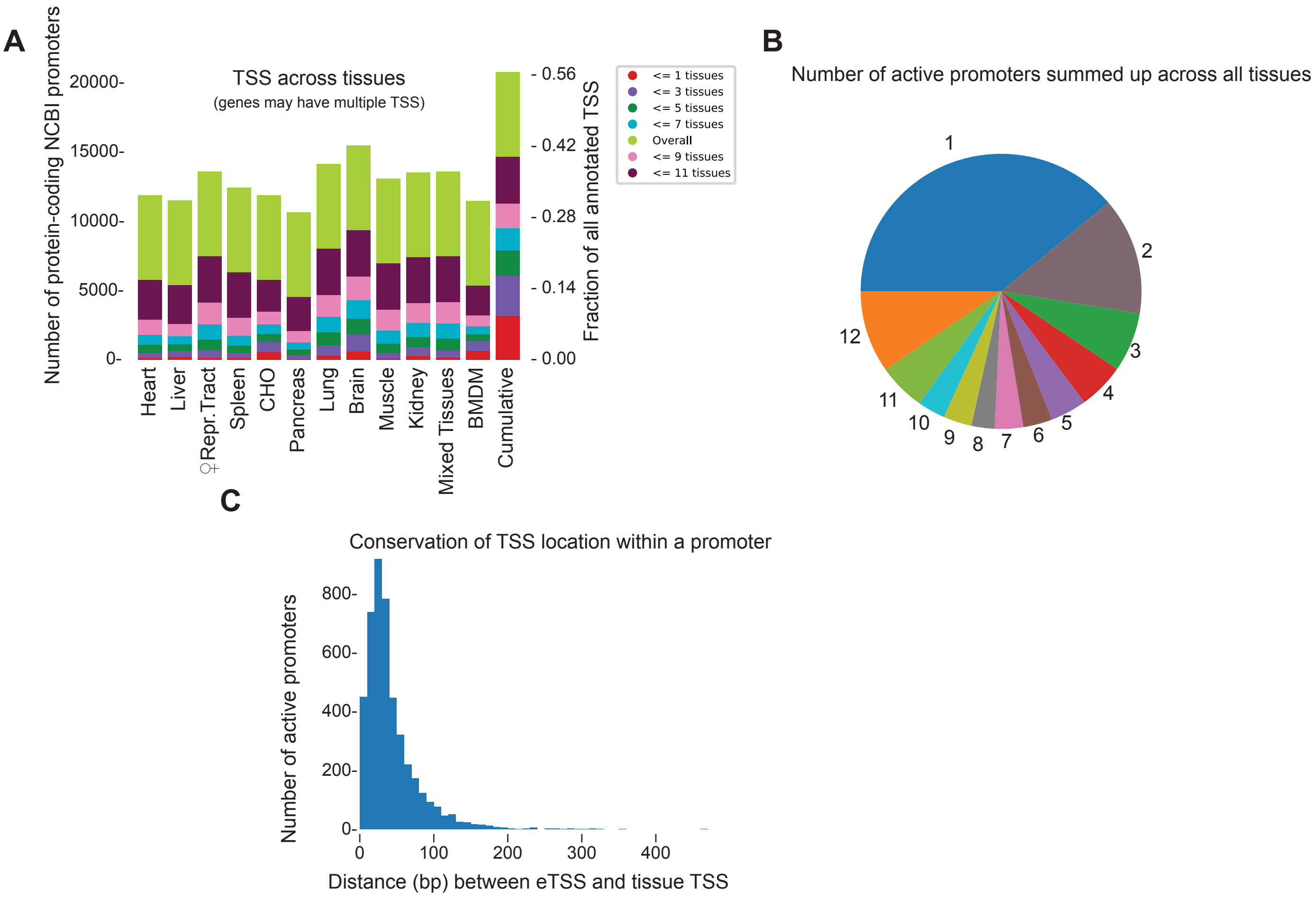


**Figure S4**

Diversity of promoter and TSS usage across tissues. (**A**) Fraction of NCBI promoters covered by each tissue and cell-type. This includes alternative start sites of isoforms. (**B**) Number of active promoters of conserved expressed genes (all tissues contain a TSS in the gene) summed up across all tissues. (**C**) Distribution of the maximum distance (bps) between the final merged eTSS output and the furthest tissue TSS in the same TSS region across conserved promoters (all tissues express the promoter).

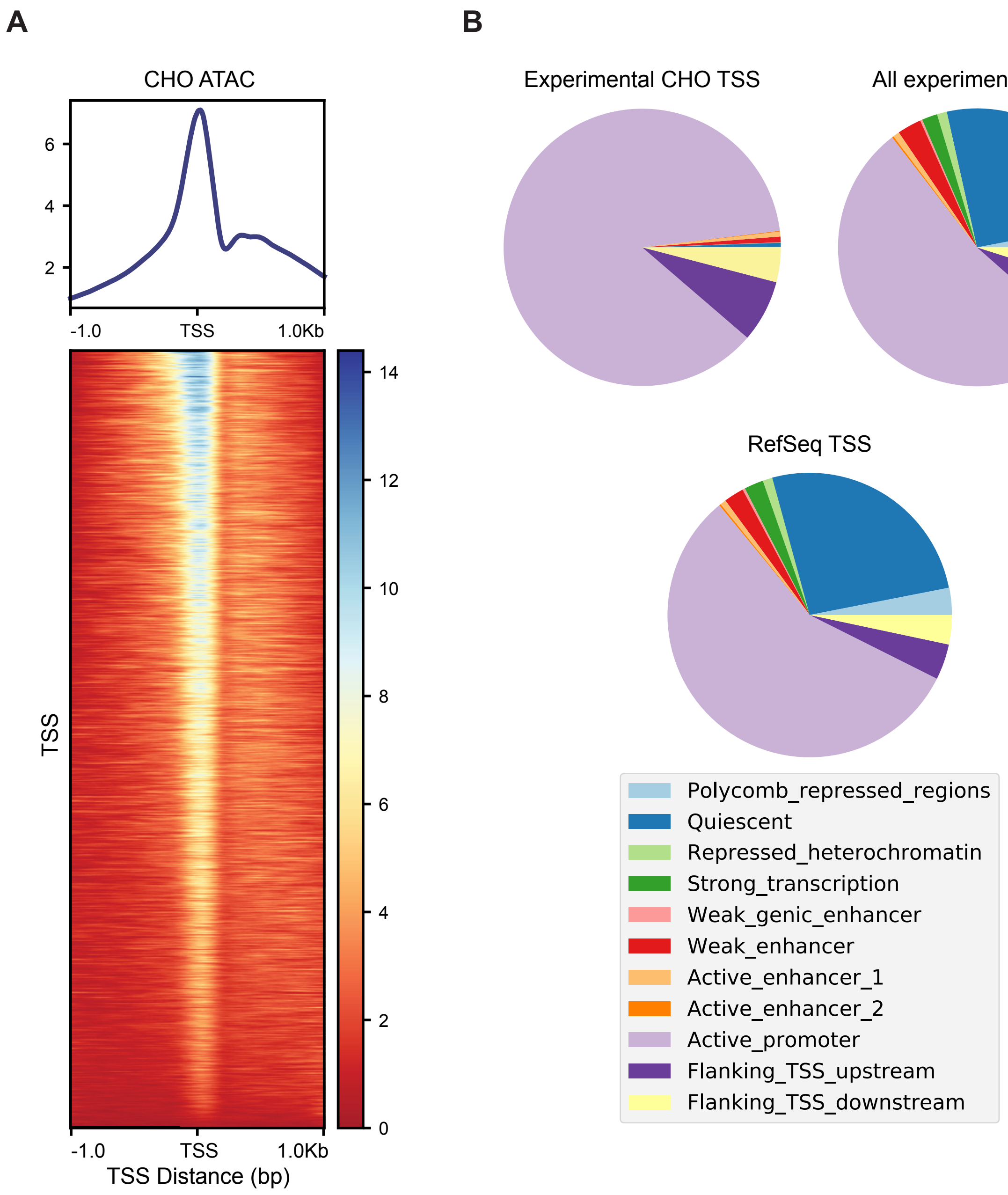


**Figure S5**
(A) ATAC-Seq pileup over TSS regions. Values are negative log10 p-value of the basepair being part of an open-chromatin peak. Top: Histogram of a CHO ATAC-Seq sample using all eTSSs that were expressed in CHO. Bottom: The same regions, in heatmap form, where each row is an eTSS. (B) Overlap of protein-coding TSS with chromatin modifications: our experimental CHO TSS, our updated annotation (all experimental TSS) and the RefSeq grouped by chromatin state made using chromHMM with histone marks from CHO in Feichtinger et al 2016 and mapped to the PICR genome.

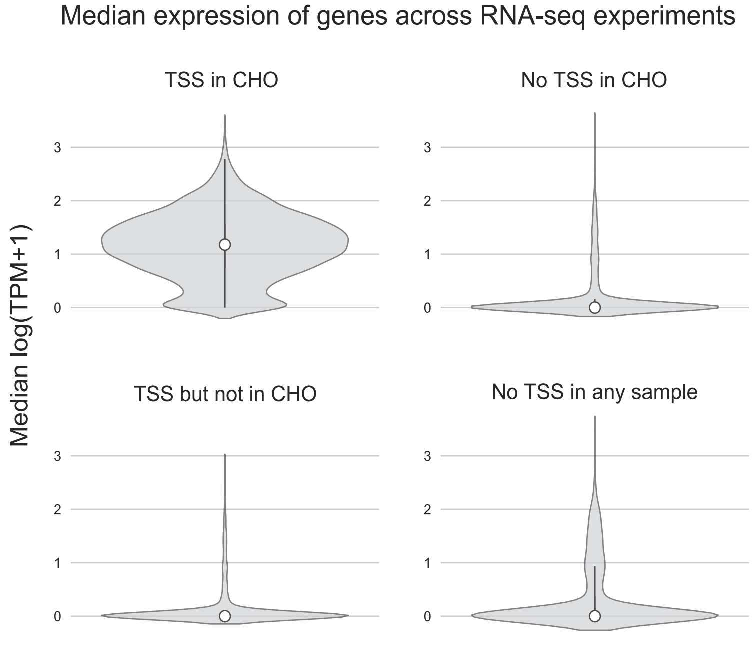


**Figure S6**
Violin plots of gene expression in CHO cells, grouped based on experimental TSS findings. 1,558 RNA-seq samples were used across different CHO cell lines and experiments. Expression is increased in genes where there are TSSs.
